# Convergent evolution associated with the loss of developmental diapause may promote extended lifespan in bees

**DOI:** 10.1101/2024.05.07.592981

**Authors:** Priscila K. F. Santos, Karen M. Kapheim

**Affiliations:** Utah State University, Logan, UT, 84322, United States of America

**Keywords:** Eusociality, comparative genomics, selection, dormancy, aging

## Abstract

Diapause has long been proposed to play a significant role in the evolution of eusociality in Hymenoptera. Recent studies have shown that shifts in the diapause stage precede social evolution in wasps and bees, however, the genomic basis remains unknown. Given the overlap in molecular pathways that regulate diapause and lifespan, we hypothesized that the evolutionary loss of developmental diapause may lead to extended lifespan among adults, which is a prerequisite for the evolution of eusociality. To test this, we compared 27 bee genomes with or without prepupal diapause. Our results point to several potential mechanisms for lifespan extension in species lacking prepupal diapause, including the loss of the growth hormone PTTH and its receptor TORSO, along with convergent selection in genes known to regulates lifespan in animals. Specifically, we observed purifying selection of pro-longevity genes and relaxed selection of anti-longevity genes within the IIS/TOR pathway in species that have lost prepupal diapause. Changes in selection pressures on this pathway may lead to the evolution of new phenotypes, such as lifespan extension and altered responses to nutritional signals, that are crucial for social evolution.

**Significance:** Diapause has long been proposed to play a significant role in the evolution of eusociality in Hymenoptera. Recent studies have shown that the loss of diapause during the prepupal stage precedes social evolution in wasps and bees. However, the genomic mechanisms underlying this phenomenon remain unknown. Through comparative genomics, we showed that the convergent loss of prepupal diapause is associated with mechanisms that may promote lifespan extension, a prerequisite for social evolution. These mechanisms include genes losses and signals of selection on genes related to aging.

## Introduction

The origin of eusociality is one of the major transitions in evolution, because it involves a shift from reproduction at the level of the individual to reproduction at the colony level (Szathmáry & Smith 1995). A typical eusocial colony of ants, bees, or wasps includes a long-lived queen who specializes on egg-laying living alongside her non-reproductive daughters who perform nest maintenance, foraging, and brood care (Wilson 1971). Understanding the selective pressures shaping this major transition requires investigating the origins of each of its defining features; (1) overlapping generations, (2) reproductive division of labor, and (3) cooperative brood care (Batra 1966; Michener 1969). For example, one of the preconditions for overlapping generations is that nest foundresses (queens) must live long enough to overlap with their adult daughters (workers). This suggests that the origins of eusociality were preceded by an extension of lifespan in the solitary ancestors (Michener 1974; Carey 2001; da Silva 2022). A potential source of this lifespan extension could be rooted in the phenomenon of diapause. Diapause is a period of dormancy in insects, in which development and reproduction are suppressed, usually as a way to pass unfavorable periods such as extreme temperatures, drought, or scarcity of resources (Denlinger 2022). In bees, diapause most commonly occurs in larvae following the last molt, a stage referred to as the prepupa (Gauld & Bolton 1988; P.K.F. Santos et al. 2019). However, there is a great deal of variation between bee species, with many bees diapausing as adults, and many skipping diapause altogether (P.K.F. Santos et al. 2019). Recent research indicates that prepupal diapause is the ancestral state in bees, but it has been lost in the ancestor of every clade in which eusociality has evolved (P.K.F. Santos et al. 2019). This pattern has been found in both bees and wasps (da Silva 2021), suggesting that while diapause itself may not be essential for social evolution, its loss during the prepupal stage could be pivotal if it leads to increased longevity, thus facilitating overlapping generations as required for the evolution of eusociality.

A recent study of lifespan evolution in Hymenoptera gleaned longevity estimates from the literature to show that primitively eusocial nest foundresses do indeed live significantly longer than non-eusocial Hymenoptera foundresses (da Silva 2022). Although diapause was not included in the final model of foundress lifespan (likely due to missing data in more than a quarter of the species), foundresses from species included in the study that have lost prepupal diapause had median lifespans 3.5 times as long as those with prepupal diapause (median = 87 d vs 25 d) (da Silva 2021). Here we attempt to explain these connections between lifespan, eusociality, and the loss of prepupal diapause by identifying molecular pathways under selection with the loss of prepupal diapause in bees.

Many of the molecular pathways that regulate diapause in insects also regulate aging and longevity (Hutfilz 2022). Although diapause at different life stages shares common features – such as reduced metabolism, increased lipid reserves, resistance to extreme environmental conditions, and prolonged lifespan – its physiological regulation varies. In prepupal diapause, development is halted due to a reduction in the production or release of the prothoracicotropic hormone (PTTH) by the prothoracic gland (PG), leading to decreased ecdysone levels, which delay metamorphosis (Denlinger 2022b; Rewitz et al. 2009). Conversely, adult diapause involves the cessation of reproduction through the arrest of oocyte development, primarily regulated by a shutdown in juvenile hormone (JH) production (Denlinger 2022b; C.G. Santos et al. 2019). Despite these differences, certain pathways regulate diapause across life stages. The insulin/insulin-like signaling and target of rapamycin (IIS/TOR) pathway is one of these pathways, known for its role in regulating both diapause and aging. The pathway’s specific regulatory functions depend on the life stage, indicating that the same pathway may experience different selective pressures depending on the stage: during developmental diapause, the IIS/TOR pathway manages nutrient allocation for growth and metamorphosis, while during adult diapause, it allocates nutrients for maintenance and reproduction (Denlinger 2022b). This suggests that shifts in the diapause stage could lead to changes in pathways regulating aging, potentially facilitating lifespan extension.

One other important difference between prepupal and adult diapause is the opportunity for cellular regeneration. Diapause during development is followed by the renewal of somatic tissues through cell divisions and the process of metamorphosis itself (Tatar & Yin 2001). Metamorphosis allows for a restructuring of the organism’s body, which can rejuvenate tissues, with potentially lifespan extending effects (Gilbert 2000; Seifert et al. 2012). However, in adults, most of the tissues are post-mitotic, meaning they cannot undergo cell division to renew themselves, limiting the extent to which lifespan can be extended by diapause alone. Adult insects that go through diapause mitigate its potential negative effects by reducing age-specific mortality through a reduced rate of senescence, an adaptation specific to the adult stage (Tatar & Yin 2001; Easwaran & Montell 2023). The loss of prepupal diapause associated with adaptations to reduce senescence rate may be key in promoting the overlap of generations and the evolution of social behavior in bees.

If shifts in diapause stage result in prolonged lifespan, then the convergent loss of developmental diapause in bees could be linked to genomic changes associated with mechanisms that regulate adult lifespan extension. To test this hypothesis, we compared the genomes of 27 bee species with and without prepupal diapause. We looked for signals of positive, purifying, and relaxation of selection, as well as gene family gain and losses linked to the absence of prepupal diapause. Next, we assessed whether genes under selection were enriched for longevity or oxidative stress-related functions. We found that the loss of prepupal diapause might be associated with mechanisms regulating adult lifespan extension through processes including the loss of the growth prothoracicotropic hormone (PTTH) and its tyrosine-protein kinase receptor torso (TORSO), purifying selection of pro-longevity genes, and relaxed selection of anti-longevity genes in the IIS/TOR pathway.

## Results

### Genomic data selection and orthogroups identification

Out of the 74 bee genomes with available annotation files and information regarding the diapause stage, 71 genomes exhibited over 80% complete BUSCO genes within their protein-coding sequences. Following the exclusion of species from the same genus sharing identical life-history traits, we retained 27 species for subsequent selection analysis (Figure 1; Table S1).

**Figure 1:**
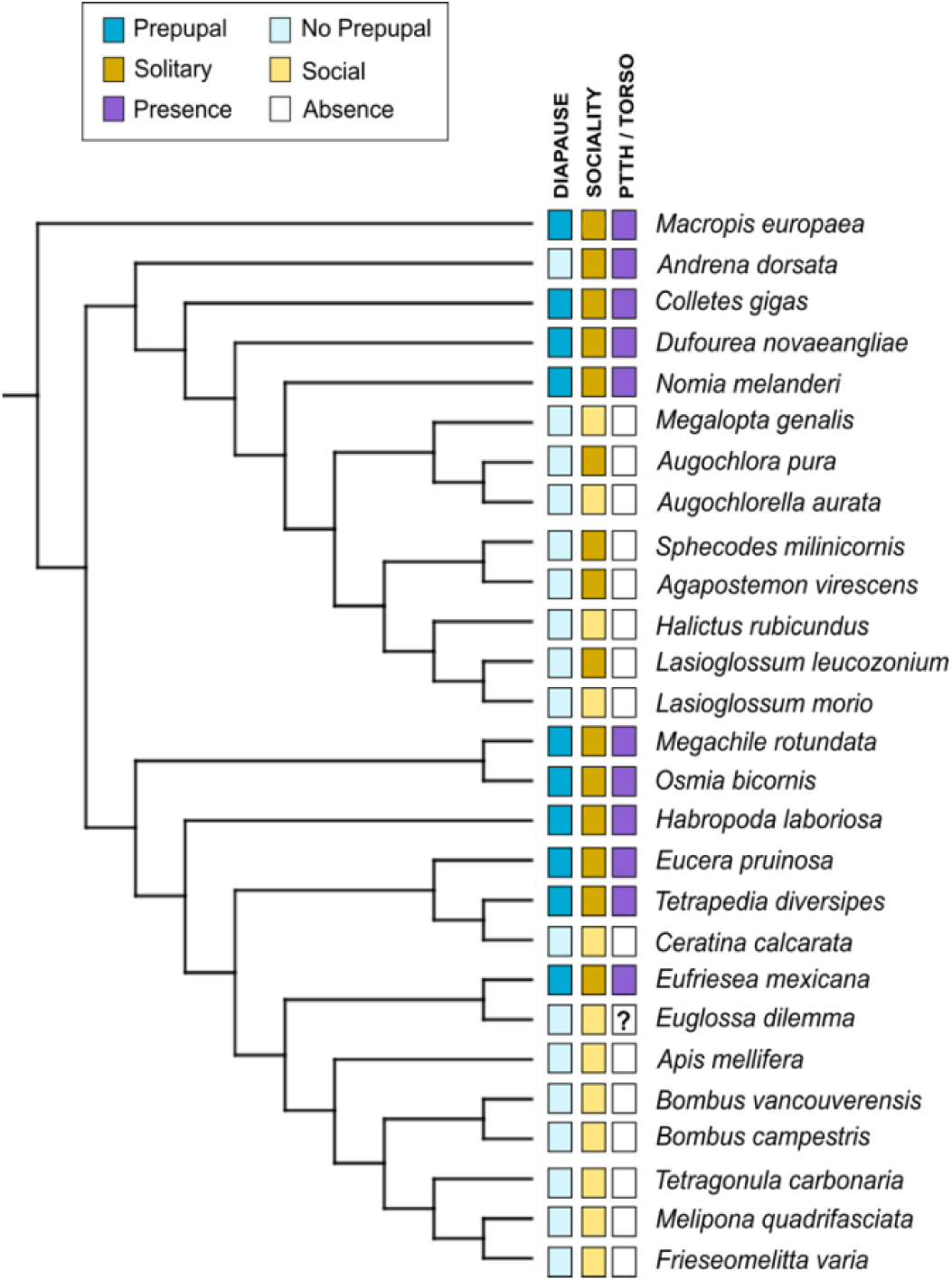
The 27 species included in the molecular evolution analysis and their associated life-history traits. The prepupal diapause stage is depicted in dark blue, while the loss of developmental diapause is represented in lighter blue. Solitary bees are indicated by a stronger yellow color, and social species are denoted by a lighter yellow. Purple represents the presence of the prothoracicotropic hormone (PTTH) and its tyrosine-protein kinase receptor torso (TORSO), while the absence of color indicates orthologs could not be found. The two species marked with “?” had conflicting results regarding the presence of the genes, depending on whether the search was conducted using the genome or the annotated protein-coding genes set as a reference.

Orthofinder analysis yielded 2592 single-copy orthogroups across all 27 species. After filtering to retain orthogroups present in all species, but being single-copy in at least 70% of them, we identified and utilized 4173 orthogroups for further analysis.

### Predominance of relaxation of selection linked to the loss of prepupal diapause

The search for convergent evolutionary rate shifts linked to the loss of prepupal diapause using RERconverge resulted in 914 significant orthogroups (Benjamini-Hochberg adjusted p-value < 0.05 after permulation). Among these orthogroups, 336 exhibited positive Rho values, signifying genes undergoing accelerated evolution (attributed to the release of selective constraint or adaptation), whereas 578 showed negative Rho values indicating a decelerated evolutionary rate (increased selective constraint) in species without diapause during the prepupal stage (Table S2).

aBSREL identified 98 orthogroups where positive selection occurred on a proportion of branches in species lacking diapause in the prepupal stage. However, the significance in most groups is associated with one species (at most three species), suggesting a weak convergent signal of positive selection related to the loss of prepupal diapause (Table S3). Using Hyphy-RELAX, 367 orthogroups were identified under intensification of selection (potentially positive or purifying selection) and 1486 orthogroups under relaxation of selection (Table S3).

Combining results from RERconverge and Hyphy, we categorized orthogroups into groups of interest (see Material and Methods for details): possible positive selection – 293 orthogroups; possible purifying selection – 347 orthogroups; possible relaxed selection – 1217 orthogroups (Figure 2A); and possible intensified selection – 260 orthogroups (Figure 2B). We also classified these orthogroups as having the best evidence of positive selection (66 orthogroups) – those that were significant in at least two of the following: aBSREL, RERconverge with Rho positive values, and intensification of selection in RELAX (Figure 2B and Table S4); or having the best evidence of purifying selection (45 orthogroups) – orthogroups significant in both RERconverge with Rho negative values and intensification of selection from RELAX (Figure 2B; Table S5).

**Figure 2:**
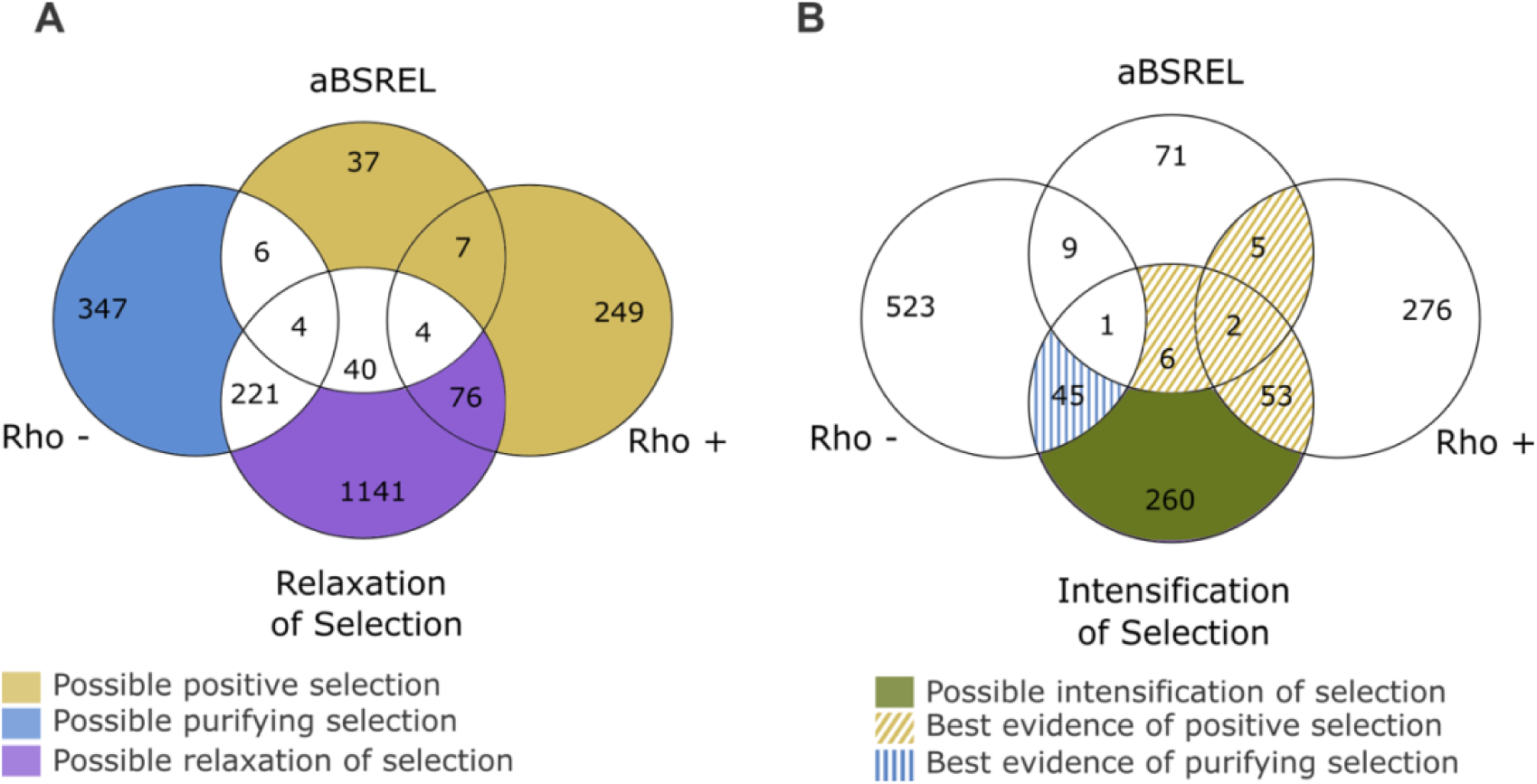
The Venn diagram depicts orthogroups categorized under relaxed (A) or intensified (B) selection, which intersect with genes under positive selection (aBSREL), accelerated evolution (Rho +), and/or decelerated evolution (Rho -). Different colors denote orthogroups potentially experiencing positive (yellow), purifying (blue), relaxed (purple), or intensified (green) selection, while dashed lines indicate orthogroups with the strongest evidence for positive (yellow) and purifying (blue) selection. Refer to the Material and Methods section for comprehensive classification details.

### Gene ontology enrichment

Orthogroups exhibiting possible positive and intensified selection were significantly enriched for terms associated with post-translational protein modifications, such as protein dephosphorylation and ubiquitin protein ligase activity, respectively. Regulation of transcription was also among the enriched terms for both positive and purifying selection (Table S6). There were no significant terms among the orthogroups under relaxed selection at p < 0.01, with double-strand break repair and RNA methylation being the top terms (p = 0.031) (Table S6).

### Prepupal diapause loss is a strong predictor of *PTTH* and *torso* loss

An analysis of gene family gain and loss revealed that only two orthogroups were lost in the majority of species that do not undergo diapause during development (Table S7). Specifically, the prothoracicotropic hormone (*PTTH*) gene and its tyrosine-protein kinase receptor torso (*torso*) were absent in all species lacking prepupae diapause, except for *Andrena dorsata*, which has both orthogroups and two copies of the *torso* receptor (Figure 1, Table S7). Our results show these losses are followed by positive selection in genes of the EGFR signaling pathway, which is the main pathway producing ecdysone during metamorphosis (Cruz et al. 2020) (Table S4). Notably, *epidermal growth factor receptor substrate 15* (*egfr15*) and *son of sevenless* (*SOS*) genes show strong evidence of positive selection (Table S4 and Figure 3).

**Figure 3:**
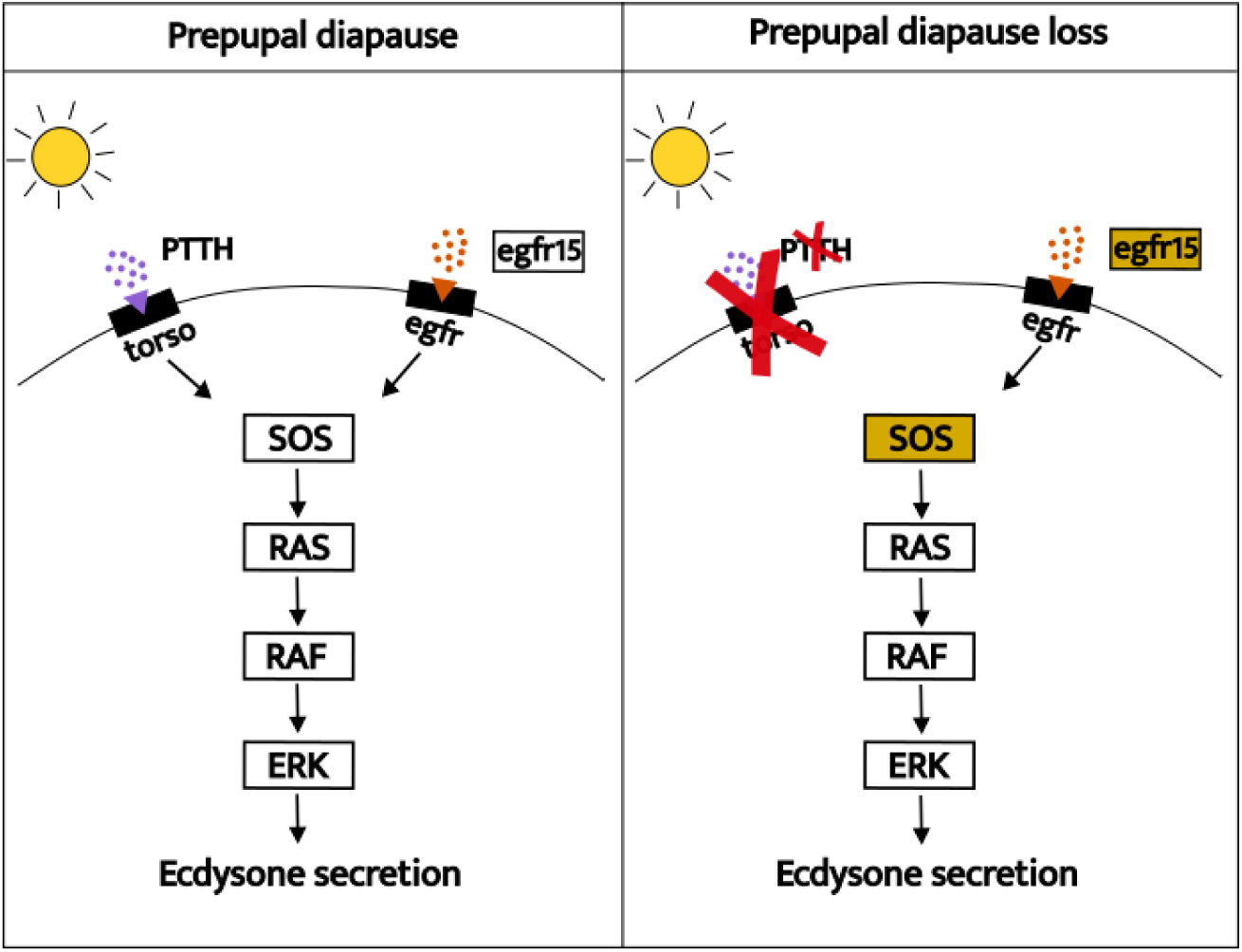
Simplified scheme illustrating two pathways regulating the production and release of ecdysone by the prothoracic gland (PG). The pathway utilizing the tyrosine-protein kinase receptor TORSO as a receptor for the prothoracicotropic hormone PTTH is absent in species that do not undergo prepupal diapause. egfr substrate 15 (EGFR15), a substrate of the alternative pathway that uses EGFR as a receptor, is subject to positive selection, along with the *son of sevenless* (*SOS*) gene, highlighting the significance of this pathway in species without prepupal diapause.

*PTTH* and *torso* loss have been previously associated with social behavior in bees. In a previous screening of 22 bee genomes, all 15 of the included social species were missing both *PTTH* and *torso* (Costa et al. 2022). We aimed to discern whether the diapause stage or sociality exerted a stronger influence on gene loss. To address this, we performed PGLS analyses, considering both predictor variables independently and simultaneously (Table S8). When analyzed individually, both diapause stage (Estimate: 0.81, Multiple R-squared: 0.87, p = 2e-13) and sociality (Estimate: 0.48, Multiple R-squared: 0.44, p=8e-05) significantly predicted the presence/absence of *PTTH* and *torso*. However, in the joint analysis, diapause stage is the only significant predictor (Estimates: Diapause(yes) – 0.81, Sociality(solitary) – 0.10; p-values: Diapause – 3e-10, Sociality – 0.1186). Furthermore, the model with both predictors had a slightly higher Multiple R-squared = 0.88 (vs 0.87 including only diapause as predictor), indicating diapause stage is a stronger predictor of *PTTH* and *torso* loss.

### Orthogroups under selection are not enriched for oxidative stress and lifespan genes

Out of 198 genes related to aging from the GenAge database (Tacutu et al. 2018), we identified 101 among our 4172 orthogroups. We found aging genes among all genes we identified as under possible positive, purifying, intensified, and relaxed selection associated with the loss of prepupal diapause, with the orthogroups under relaxation of selection including the highest number of aging genes (26 out of a total of 1217 genes). However, none of the groups exhibited more aging genes than expected by chance (Hypergeometric test: p > 0.2). Many of the aging genes under selection are part of the IIS/TOR pathway (Figure 4). One gene previously described as pro-longevity, and at least five genes described as anti-longevity in *Drosophila* (Tacutu et al. 2018) were identified as under purified and relaxed selection in the current study, respectively (Figure 4 and Table 1).

**Figure 4:**
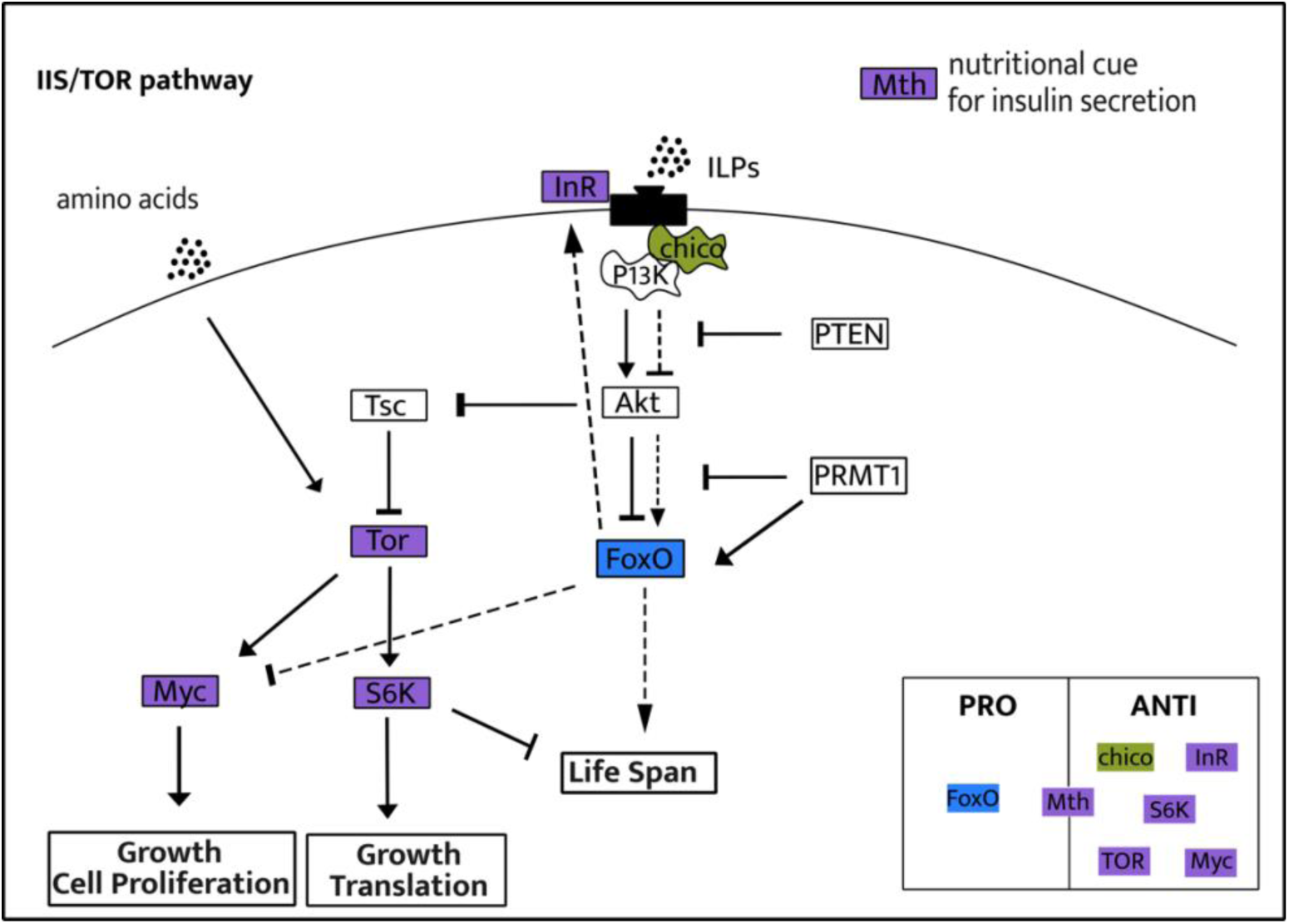
Simplified scheme of the insulin/insulin-like signaling (IIS) and target of rapamycin (TOR) pathways containing genes under selection in the current study. When the insulin pathway is activated (high nutrition), Akt phosphorylates FoxO, preventing its entrance to the nucleus of the cell, and activates Tor, leading to growth. When there is a reduction in the IIS/TOR pathway activation (low-nutrition), FoxO enters in the nucleus of the cells, regulating thousands of genes, leading to lifespan extension. Continuous arrows represent activation of the pathway in an environment with high nutrition, for example, while dashed arrows represent the alternative pathway in environments with low nutrition. Genes within green, blue, or purple boxes represent those under intensified, purifying, and relaxed selection in the current study, respectively. The box in the bottom right classifies the genes as pro- or anti-longevity based on their known effects on *Drosophila* lifespan (Tacutu et al. 2018). Figure made based on Kapahi and Zid 2004; Dutriaux et al. 2013; Texada et al. 2020; Denlinger 2022.

**Table 1:**
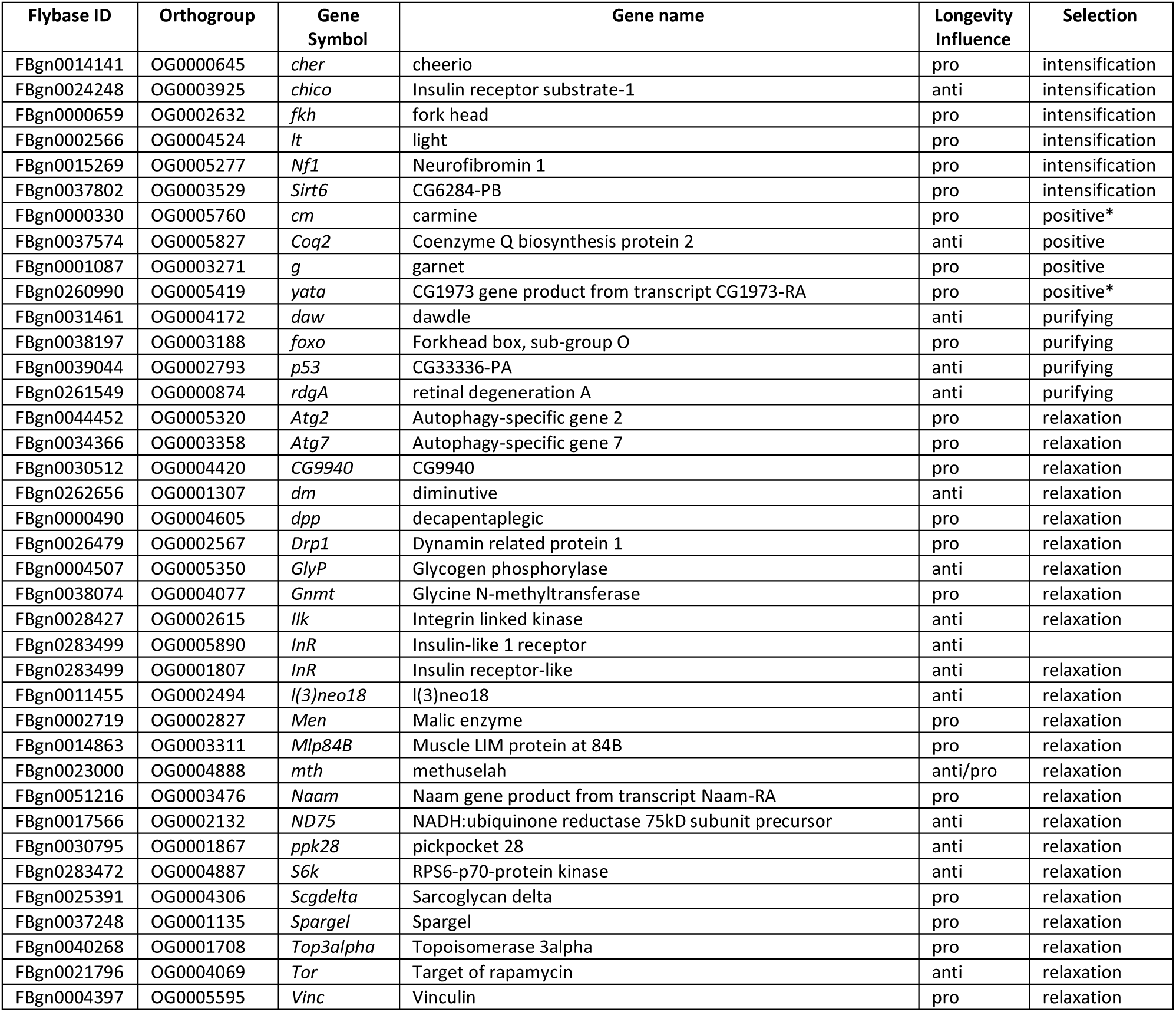
Orthogroups under selection overlapping with the *Drosophila* aging genes list from GenAge database (Tacutu et al. 2018). Longevity influence classifies the orthologs as pro- or anti-longevity genes according to their effects on *Drosophila* lifespan. The asterisk (*) highlights orthogroups that are also part of the best evidence of positive selection (Table 1).

Regarding oxidative stress genes (Kramer et al. 2021), 17 out of 34 had orthologs among the bee orthogroups. Only the positive selection and the relaxation of selection groups included oxidative stress orthologs in their lists, although not exceeding the amount of overlap expected by chance (Hypergeometric test: p > 0.7).

## Discussion

In this study, we tested the hypothesis that the evolutionary loss of prepupal diapause is followed by genomic changes that might be associated with adult lifespan extension, a prerequisite for the evolution of sociality in bees (Batra 1966; Carey 2001; Michener 1974). Our findings revealed convergent signals of selection linked to the loss of prepupal diapause, with the majority of genes experiencing relaxed selection. However, there is no significant overlap between genes under selection and genes previously associated with aging or oxidative stress in other groups. Below we discuss specific pathways that may be relevant to the adult lifespan extension in the absence of prepupal diapause. These include endocrine pathways that regulate metamorphosis, the IIS/TOR lifespan regulator pathway, and genes related to cell cycle and DNA repair damage.

### Loss of *PTTH/torso* promotes selection in the EGFR pathway

The loss of *PTTH and torso* along with positive selection in the *egfr15* and *SOS* genes (Figure 3) suggest a change in the dynamics of the metamorphosis pathways when prepupal diapause is lost. The onset of metamorphosis in holometabolous insects is regulated by the production and release of ecdysone by the prothoracic gland (PG) in the brain (Henrich et al. 1999). When PTTH binds to its tyrosine kinase receptor TORSO in the PG, the Ras/RAF/Erk pathway is activated, which stimulates the release of ecdysone (Pan & O’Connor 2021). Although unknown for bees, in other insects PTTH also plays a pivotal role in larval and pupal diapause. During diapause, the brain-PG axis is shut down, inhibiting PTTH production and release, subsequently reducing production of ecdysteroids, resulting in delayed metamorphosis (Figure 3) (Denlinger 2022).

For a considerable period, PTTH/TORSO was considered the primary pathway for ecdysone production through the activation of the PG (Rewitz et al. 2009). However, there are at least four other tyrosine kinase receptors expressed in the PG. A recent study demonstrated that most of these are dispensable, because EGFR is the only receptor whose absence prevents metamorphosis in flies (Cruz et al. 2020). This suggests EGFR is the ancestral ecdysone biosynthesis pathway, with PTTH/TORSO likely co-opted in holometabolous insects to finely adjust the timing of pupation in response to environmental cues (Shimell et al. 2018; Cruz et al. 2020). Recurrent *PTTH* and *torso* losses across species support the hypothesis that EGFR is the foundational pathway regulating metamorphosis (Cruz et al. 2020). Moreover, it suggests that PTTH/TORSO are dynamic elements susceptible to environmental influences (Skelly et al. 2019). Evidence of positive selection in two genes of the EGFR pathway indicates an amplified role of this pathway in the absence of *PTTH* and *torso* in bees (Figure 3).

In Hymenoptera, the loss of *PTTH* and *torso* appears linked to the evolution of sociality in bees, as all examined social bee genomes lack these genes (Costa et al. 2022) (Figure 1). Our findings suggest that not only sociality, but also the loss of developmental diapause, predicts *PTTH* and *torso* loss, with the latter being a stronger predictor. Since the loss of developmental diapause is also a strong predictor of sociality in bees, future analysis of additional bee genomes will be necessary to fully separate these effects.

The near-universal loss of *PTTH* and *torso* only in species lacking prepupal diapause suggests strong selection against this pathway in the species that do not go through diapause during development. In *Drosophila*, reducing expression of *PTTH* or *torso* delays pupariation (Shimell et al. 2018; Kang et al. 2023; Rewitz et al. 2009), while constitutively knocking down *PTTH* or *torso* in the PG results in prolonged adult lifespan, suggesting possible detrimental effects of *PTTH* and *torso* in adulthood (Kang et al. 2023). This aligns with the antagonist pleiotropy theory of aging (Williams 1957), where genes like *PTTH* and *torso* that confer benefits early in life (e.g. promoting survival via diapause induction), but with detrimental effects later in life (reduction of adult lifespan), are favored by natural selection and contribute to aging.

We propose that *PTTH* and *torso* loss is a consequence of the loss of prepupal diapause and precedes the evolution of sociality. Selection may act against the PTTH/TORSO pathway once it is no longer needed for diapause functions due to its lifespan reducing effects in adults. In this way, the adult lifespan extension requisite for eusociality may be a result of reduced antagonistic pleiotropy, freeing PTTH/TORSO from the selection shadow once their function in early life was eliminated.

### Loss of developmental diapause relaxes selection in the IIS/TOR pathways

Several genes in the insulin/insulin-like signaling (IIS) and target of rapamycin (TOR) pathways are under relaxed selection in species that have lost prepupal diapause (Figure 4). These pathways are highly conserved among metazoans and are key regulators of aging (Bitto et al. 2015; Guarente & Kenyon 2000). IIS and TOR work in parallel and interactively, such that when either or both are downregulated, there is an increase in lifespan (Kapahi & Zid 2004; McCormick et al. 2011).

In our study, *FoxO*, a pro-longevity gene in *Drosophila*, is under purifying selection in bee species that lost prepupal diapause, reinforcing the importance of this gene, independent of the diapause stage. All the anti-longevity genes in this pathway are under relaxed selection (Figure 4). The exception is *mth*, which has both pro- and anti-longevity properties. Relaxed selection in genes of the IIS/TOR pathway has been associated with both reduction of lifespan, as observed in annual vs long-lived killifish (Cui et al. 2019), and increased lifespan as seen in long vs short-lived mammals (Liu et al. 2023). Moreover, aging genes are under relaxed selection in humans compared to chimpanzees, despite humans living approximately twice as long (Cui et al. 2019). Similarly, relaxed selection on genes in the IIS/TOR pathway in bees that have lost prepupal diapause could be associated with increased lifespan.

The hyperfunction theory of aging predicts that reduction in lifespan results from the inappropriate activation of IIS/TOR pathway when growth is no longer beneficial (Blagosklonny 2006; Gems 2022). In accordance with this hypothesis, relaxed selection in IIS/TOR genes could be associated with loss or shifts in function in some of the genes in this pathway, which could result in reduced activation and increased lifespan. However, IIS/TOR is an essential pathway for holometabolous development, so a complete loss of function is unlikely (Dutriaux et al. 2013; Fernandez et al. 1995; Hansen et al. 2004). Relaxed selection has also been associated with the evolution of new phenotypes through the release of constraint in traits that were under opposing selection pressures in different environments (Hunt et al. 2011). In Hymenoptera, relaxed selection of caste-biased genes preceded the evolution of sociality and could be the basis for phenotypic plasticity. Moreover, in honeybees, fast evolving genes were more likely to be differentially expressed between castes (Hunt et al. 2011).

Loss of constraint in the IIS/TOR pathway suggests that different responses to nutritional cues could evolve, a hallmark of caste differentiation in many social species (Kapheim 2017). Differences in diet, either qualitative or quantitative, are associated at some level with caste determination in all independent origins of castes in bees (Starr 2020; Carnell et al. 2020; Kapheim et al. 2011; Richards & Packer 1994; Rehan & Richards 2010). A notable instance of a unique nutritional response is observed in honeybee queens. Consumption of royal jelly, a highly nutritious food, by the queens, leads to decreased activation of insulin pathway, resulting in lifespan extension and increased fertility (Corona et al 2007). This is the opposite effect of what is expected to other animals, in which a reduction of activity in the insulin pathway leads to reduced lifespan and fertility (Ament et al. 2008). Relaxed selection on genes in the IIS/TOR pathway could allow for such reversals to evolve.

Interestingly, a link between nutrition, caste differentiation, and diapause has long been recognized (Hunt and Amdam 2005; Hunt 2006; Hunt et al. 2007). Our results suggest a possible mechanism for this connection. Specifically, shifts in the timing of diapause may have allowed molecular pathways important during development, such as IIS/TOR, to become co-opted for social evolution (Hunt and Amdam 2005). We suggest that relaxation of the IIS/TOR pathway, as consequence of the loss of prepupal diapause, precedes the evolution of sociality and could be related to lifespan extension and/or the evolution of new phenotypes in response to differences in nutrition.

### Many cell cycle regulation and DNA damage repair genes are under positive selection in species lacking prepupal diapause

Many genes involved in cell cycle regulation and DNA damage repair are subject to selection in bee species that have lost prepupal diapause. These two biological processes are crucial for diapause and aging mechanisms.

Diapause in the adult stage may require specific adaptations to maintain cell cycle arrest without accumulating damage, given the lack of somatic tissue renewal in adults of holometabolous insects (Tatar & Yin 2001). For instance, Fizzy (*fzr*), which is under positive selection in bee species lacking prepupal diapause (Table S4), is involved in the natural response to injuries in adult flies. Endocycles (consecutive cycles of cell growth and arrest in the M phase), rather than mitosis, is the default response to injury in adult flies (Cohen et al. 2018). However, the expression of *fzr* alone is sufficient to override the endocycle response, activating mitosis after injury instead (Cohen et al. 2018). Positive selection of the *fzr* gene in bees that have lost prepupal diapause suggests a potential mechanism to prevent organ injuries under adverse environmental conditions, which could indirectly extend adult lifespan.

Other genes involved in cell cycle regulation and cell proliferation control, such as tumor suppressor genes, are under positive selection in species that have lost prepupal diapause (Table S4). Examples include *G1/S-specific cyclin-E1* and *cyclin-dependent kinase 14*, which regulates the transition between phases G1 and S of the cell cycle (Chen et al. 2019; Fagundes & Teixeira 2021), *ST7* and *TRIM33* that act as tumor suppressor genes (Charong et al. 2011; Xue et al. 2015), and *Kruppel-like factor 6,* a transcription factor involved in the progression of multiple malignant tumors (Sabatino et al. 2019). *p53* (Table 1), which also regulates cell cycle arrest at both G1/S and G2/M checkpoints (Engeland 2018), is under purifying selection. However, it remains unclear how selection on these genes could promote lifespan extension.

DNA repair mechanisms are crucial in preventing aging. Genes like *INO80 complex subunit D*, which is involved in DNA recombination and DNA repair (Poli et al. 2017; Horigome et al. 2014), and *ASH2*, which acts upstream of cellular response to DNA damage stimulus (Burgess et al. 2014), are under positive selection and are potentially adaptive for lifespan extension (Table S4). Additionally, the gene ontology term related to double-strand break repair (p = 0.031, Table S6) is enriched among the genes under relaxation of selection, suggesting loss of constraint and possible evolution of new phenotypes associated with DNA repair in those bee species.

## Conclusions

Our study supports the hypothesis that convergent evolutionary changes associated with the loss of prepupal diapause is related to molecular pathways that might regulate adult lifespan extension. Our findings indicate that this extension appears to stem from a combination of factors, such as the loss of *PTTH* and *torso*, coupled with a relaxation of selection in the IIS/TOR pathway. This relaxation of selection, due to the loss of prepupal diapause, may facilitate the emergence of new phenotypes, such as increased lifespan or altered responses to nutritional signals, two important adaptations in the evolution of castes in social insects. Given the strong correlation between loss of prepupal diapause and the evolution of social behavior in bees, disentangling the effects of each phenotype on genome evolution proves challenging.

Therefore, advancing our understanding of the link between diapause and the evolution of sociality demands expanding our understanding of the natural history of species presenting different life history traits. Solitary bee species lacking prepupal diapause offer ideal opportunities for investigating lifespan extension, examining signs of relaxed selection in *PTTH* and *torso* (if these genes are still present) compared to solitary species retaining prepupal diapause, and exploring potential innovations in how IIS/TOR pathway responds to diverse stimuli.

## Material and Methods

### Genomes

For the comparative genomics analysis, we obtained bee genomes from the National Center for Biotechnology Information (NCBI) (https://www.ncbi.nlm.nih.gov/datasets), the National Genomics Data Center-China National Center for Bioinformation (NGDC-CNCB) (https://ngdc.cncb.ac.cn/), and the Halictid Genome Browser (https://beenomes.princeton.edu). Specifically, we selected genomes for which annotation files were accessible and for which we possessed information regarding the diapause stage and the social level of the bees (Table S1). Solitary and communal species were classified as ’solitary,’ while subsocial, primitively social, and advanced eusocial species were classified as ’social.’ Species that undergo diapause in the last larval instar before pupation were categorized as having ’prepupal’ diapause, whereas species with adult, reproductive, or no diapause were classified as ’no prepupal’.

With the genome and annotation files, we used the agat-v0.9.2 software (Dainat 2023) to retrieve the longest isoform for each gene. Subsequently, based on the longest isoform, we extracted and translated the nucleotide coding sequences (CDS) for each gene. The protein coding sequences were then used to run BUSCO-v5.3.2 (Simao et al. 2015) against the Hymenoptera database, which encompasses 5991 genes. To ensure data quality, we retained genomes in which > 80% of the proteins were classified as complete by BUSCO.

In the genome selection process, we retained one genome per genus, opting for the genome with the highest percentage of complete proteins compared to the BUSCO database. More than one genome per genus was kept in cases where distinct life history traits, such as solitary versus social versus parasitic lifestyle, or variations in the diapause stage (prepupa versus adult) were evident. Ultimately, a total of 27 genomes were included in the subsequent analysis (Figure 1).

### Identification of Orthogroups

The protein coding sequences from the 27 different species served as input for Orthofinder-v2.5.4 (Emms & Kelly 2015) to classify orthogroups. To proceed with further analysis, single-copy orthologs were necessary. Orthofinder identified 2592 orthogroups with single-copy orthologues. To augment the number of orthogroups for subsequent analysis, we filtered the “Orthogroups.GeneCounts” table (output from Orthofinder) to retain orthogroups present in all species, but single-copy in at least 70% of those species. This process yielded 4173 orthogroups. In instances where species had paralogs, all copies were excluded from the orthogroups before conducting the molecular evolution analysis, resulting in some orthogroups containing fewer than 27 species.

### Orthogroups nucleotide alignments and trees

We compiled the nucleotide coding sequences of all species into a single file to construct a database. Utilizing the IDs of each sequence belonging to an orthogroup (generated using the protein sequences), we retrieved the nucleotide sequences by comparing them with the database. Species with more than one orthologous sequence (paralogs) in a given orthogroup were excluded. Subsequently, we employed the orthogroup nucleotide sequences as input for the MACSE_ALFIX_v01.sif pipeline, which encompasses codon alignment (Ranwez et al. 2021,https://hal.science/hal-03099847/document). The alignment pipeline included several steps to ensure high-quality alignments. First, a pre-filtering step was performed to mask potential untranslated regions (UTRs) and long non-homologous fragments. Next, HMMCleaner was used to filter and mask amino acid residues that appeared to be misaligned. In the post-processing stage, isolated codons were masked, and sequences that were more than 80% masked were removed. Finally, the extremities of the alignments were trimmed to ensure that at least 70% of the nucleotides (excluding gaps and ambiguous bases) were present at the first and last sites. The orthogroup OG0003883 was excluded from the subsequent analysis due to its alignment never concluding.

The resulting nucleotide alignment underwent filtering by removing positions with excessive ambiguous data using the pxclsq function from the phyx-v1.3 software (Brown et al. 2017). Specifically, alignment columns missing information for over 50% of the species were eliminated. The filtered alignment was then used as input for iqtree2-v2.2.0 (Minh et al. 2020; Kalyaanamoorthy et al. 2017) to generate phylogenetic trees for each orthogroup with the options -st CODON and -m MFP, allowing for the automatic identification of the best codon model for each alignment.

### Molecular evolution analysis

RERconverge-v0.3.0 (Kowalczyk et al. 2019) was used to compute convergent evolutionary rate shifts associated with the loss of developmental diapause. The input comprised nucleotide alignments from MACSE and the species tree generated in Orthofinder. Some branches in the species tree were adjusted using Mesquite-v3.81 (Maddison & Maddison 2023) to align with the topology presented in (Almeida et al. 2023). The ‘estimatePhangornTreeAll’ function with the codon-based model YN98 (submodel = “YN98”, type = “CODON”) was employed to predict the orthogroups tree while maintaining the same topology as the species tree. Four orthogroups were excluded for the RERconverge analysis because they exceeded the cluster’s time limit for the branch length estimation step. Seventeen species that had lost prepupal diapause (Figure 1) were used as foreground branches. Correlations were computed using the terminal clades with unidirectional transitions.

To empirically calibrate the p-values, a permulation analysis was conducted (Saputra et al. 2021), as outlined in https://rdrr.io/github/nclark-lab/RERconverge/f/vignettes/PermulationWalkthrough.Rmd. Briefly, 1000 permulated binary trait trees were generated for each orthogroup using the Complete Case (CC) method with *Macropis europaea* as the root species. The functions ‘getPermsBinary’ and ‘permpvalcor’ were used to produce null p-values and correlation statistics. Empirical p-values were calculated as the proportion of null statistics that were as extreme or more extreme than the observed parametric statistics, followed by Benjamini-Hochberg correction. The analysis was performed without specifying sister clades or including common ancestors. Orthogroups exhibiting significant shifts in evolutionary rates were identified based on corrected permulation p-values < 0.05. Orthogroups with Rho negative (Rho-) values indicate a slower evolutionary rate in the foreground species, possibly reflecting increased selective constraint. Conversely, Rho positive (Rho+) values denote genes undergoing faster evolution, which could result from a release of selective constraint or adaptation. For additional details on the RERconverge pipeline, refer to the GitHub repository.

To ascertain whether orthogroups exhibiting Rho+ values indicate an accelerated evolutionary rate due to positive selection or relaxation of selection, we ran Hyphy-v2.5.55 (Kosakovsky Pond et al. 2020) aBSREL (Smith et al. 2015) and RELAX (Wertheim et al. 2015). aBSREL identifies signals of positive selection in single branches among the tested branches, whereas RELAX discerns signals of convergent intensification of selection (K > 1, representing positive or purifying selection) and relaxation of selection (K < 1) in tested (foreground) branches compared to background branches. For both aBSREL and RELAX analyses, we used the nucleotide alignment from MACSE and the corresponding trees from IQ-TREE as input files. Test branches were specified as species that have lost developmental diapause. The parameters -- srv (yes) and --multiple-hits (Double+Triple) were employed to allow site-to-site synonymous rate variation and multiple simultaneous hits. The threshold for positive selection in single branches or convergent intensification or relaxation of selection in the tested branches was p = 0.05. aBSREL p-values was corrected by the Holm-Bonferroni method within Hyphy, and RELAX p-values were corrected using the p.adjust function in the stats R package (R Core Team 2023) using the Benjamini & Hochberg method.

### Gene ontology enrichment

To identify gene ontology enrichment in orthogroups associated with the loss of developmental diapause, we categorized the orthogroups of interest into four groups: possible positive selection, possible purifying selection, possible relaxation of selection, and possible intensification of selection. Possible positive selection included significant orthogroups with Rho+ values in RERconverge and significant orthogroups from aBSREL that did not overlap with relaxation of selection. Possible purifying selection included significant orthogroups with Rho-values from RERconverge that did not overlap with aBSREL or relaxation of selection.

Orthogroups possibly under relaxation of selection were those significant in RELAX (K<1) that did not overlap with aBSREL or RERconverge (Rho+ and Rho-) (Figure 2A). Lastly, orthogroups under intensification of selection were those significant in RELAX (K>1) that did not overlap with aBSREL or RERconverge (Rho+ and Rho-) (Figure 2B). The latter was considered a separate category because it is not possible to differentiate between positive and purifying selection.

Gene ontology (GO) terms were annotated for the 4173 orthogroups using InterProScan-v5.63 (Jones et al. 2014) and the databases CDD, Pfam, SUPERFAMILY, and PANTHER. Out of these, 3267 orthogroups had at least one GO term annotated, forming the gene universe for the enrichment analysis. We used topGO-v2.50.0 (Alexa & Rahnenfuhrer 2023) with the weight01 algorithm and Fisher’s exact test statistics to identify enriched terms in each one of the previously described categories. Terms were considered significant if they had a p-value < 0.01 and Annotated genes > 5.

### Gene family evolution

To assess changes in gene families associated with the loss of developmental diapause, we used the Orthogroup.GeneCount.tsv table generated in Orthofinder as input. The ‘clade_and_size_filter.py’ script from CAFE-v5.0 (Mendes et al. 2020) was used to filter out gene families with 100 or more genes. The Base_count.tab output table was then used to identify orthogroups that were gained or lost based on the diapause stage (prepupae diapause or loss of developmental diapause). Filtering was performed for all species collectively and for specific bee families with more than one species in each category (Apidae and Halictidae) (Table S7) using a custom bash script.

### Phylogenetic Generalized Least Squares (PGLS) analysis

We conducted a PGLS analysis using the R packages ape-v5.7.1 (Paradis & Schliep 2019) and caper-v1.0.3 (Orme et al. 2023), to test whether the diapause stage and sociality are significant predictors for the presence or absence of the prothoracicotropic hormone (*PTTH*) gene and its receptor tyrosine-protein kinase receptor torso (*torso*). The dataset for this analysis included the 27 species used in the molecular evolution analysis (Figure 1), along with *Exoneura robusta* and *Exoneurella tridentata*. The phylogeny, generated by Orthofinder, encompasses all species, and some branches were adjusted using Mesquite-v3.81 to align with the topology presented in (Almeida et al. 2023). Additionally, a table of traits, including diapause stage, sociality, and presence/absence of PTTH/torso, was utilized (Table S8).

The presence of *PTTH* and *torso* was first confirmed in the orthogroup files comprising the 29 species. Following this, a BLAST search was conducted using all protein coding sequence copies of *PTTH* from the same orthogroup (11 sequences). This search was performed against the genome of 118 species using tblastn and against the protein database of 48 species using blastp (Table S9). Although additional species were included in this search, the discussion and PGLS analysis were limited to the 29 species mentioned at the beginning of this section. This limitation was due to a lack of information on diapause stages, lack of annotation for protein retrieval and consequently the built of the phylogenetic tree, or because the species were from already represented genus, leading to redundant information.

For both BLAST searches, the ‘max_target_seqs’ was set to 100 and ‘evaluè to 1e-10. The search for *torso* resulted in multiple hits across different genome locations, making it challenging to identify the true ortholog accurately. Therefore, the presence of *torso* was only verified in the orthogroup file. Since all species that lost *PTTH* also lost *torso*, we treated them as a single trait in the PGLS analysis (Table S9). To model these traits, we used function pgls from the R package caper-v1.0.3. Because diapause stage and sociality are correlated variables (Fisher’s exact test; p=0.0001), we employed the model with each variable one at a time (PTTH.TORSO ∼ Prepupae_Diapause; and PTTH.TORSO ∼ Sociality) or both together (PTTH.TORSO ∼ Prepupae_Diapause + Sociality). The presence/absence of *PTTH* and *torso* was set as a binary trait, with 0 indicating the absence and 1 indicating the presence of the orthogroups. Diapause stage and sociality were set as categorical variables, yes/no for the presence or absence or prepupal diapause, and solitary/social for social status.

### Aging/longevity and oxidative genes

We compared the genes under selection with genes previously associated with aging/longevity or oxidative stress in other species. For aging-related genes, we used the list of the *Drosophila* aging-related genes obtained from the GenAge database (accessed in January 2024: https://genomics.senescence.info/genes/index.html, (Tacutu et al. 2018)). We utilized Entrez IDs to retrieve equivalent gene IDs and CDS nucleotide sequences from FlyBase (https://flybase.org, Gramates et al. 2022). Concerning oxidative stress genes, we used as reference the list of genes present in at least one of four Hymenoptera species from (Kramer et al. 2021) and downloaded the sequences from Flybase based on the gene name.

For both sets of genes, we performed a reciprocal blast search against the translated nucleotide coding sequences (CDS) of all 27 species combined in one file (322804 sequences). Initially, a tblastn was conducted using either the aging or oxidative stress genes as the database and the translated CDS sequences as query. Subsequently, using blastx, we used all translated CDS sequences as the database and either the aging or oxidative stress genes as queries. In all four searches ‘max_target_seqs’ was set to 1 and ‘evaluè to 1e-10. The reciprocal best hits were identified using the searchRBH.sh script adapted from https://morphoscape.wordpress.com/2020/08/18/reciprocal-best-hits-blast-rbhb/. In the final table, we included both orthogroups IDs and gene IDs based on the sequence IDs from the fasta files using a custom script. The orthogroup OG0005890, which is the best orthologous match to the *Drosophila insulin receptor* gene (InR), was initially excluded from the analysis due to its absence in one species. However, since OG0005890 also corresponds to an InR, it has been included in Table 1.

To assess whether any of our gene groups under selection (positive, purifying, relaxation, or intensification) were enriched for aging or oxidative stress genes, we used the phyper function from the hypeR-v2.0.0 R package (Federico & Monti 2020).

## Acknowledgments

The support and resources from the Center for High Performance Computing at the University of Utah are gratefully acknowledged. Special thanks to Tim Delory for support with snakemake and discussions of the data analysis pipeline. We are grateful to the Kapheim Lab group for their comments on earlier versions of the manuscript.

## Data availability

All scripts, input, and output files used in the analyses are available at: https://github.com/kapheimlab/Comparative_genomics_loss_prepupal_diapause. Hyphy jsonfiles will be available upon request.

